# Scale-specific analysis of fMRI data on the irregular cortical surface

**DOI:** 10.1101/224873

**Authors:** Yi Chen, Radoslaw Martin Cichy, Wilhelm Stannat, John-Dylan Haynes

**Affiliations:** Bernstein Center for Computational Neuroscience, Berlin Center of Advanced Neuroimaging & Clinic of Neurology, Charité–Universitätsmedizin Berlin, corporate member of Humboldt-Universität zu Berlin, Freie Universität Berlin, and Berlin Institute of Health, Berlin, Germany; Institute of Cognitive Neurology and Dementia Research, University Hospital Magdeburg, Magdeburg, Germany; Department of Education and Psychology, Free University Berlin, Berlin, Germany; Institute for Mathematics, Technical University Berlin, Berlin, Germany

## Abstract

To fully characterize the activity patterns on the cerebral cortex as measured with fMRI, the spatial scale of the patterns must be ascertained. Here we address this problem by constructing steerable bandpass filters on the discrete, irregular cortical mesh, using an improved Gaussian smoothing in combination with differential operators of directional derivatives. We demonstrate the utility of the algorithm in two ways. First, using modelling we show that our algorithm yields superior results in numerical precision and spatial uniformity of filter kernels compared to the most widely adopted approach for cortical smoothing. An important interim insight hereby was that the effective scales of information differ from the nominal filter sizes applied to extract them, and thus need to be calculated separately to compare different algorithms on par. Second, we applied the algorithm to an fMRI dataset to assess the scale and pattern form of cortical encoding of information about visual objects in the ventral visual pathway. We found that filtering by our method improved the detection of discriminant information about experimental conditions over previous methods, that the level of categorization (subordinate versus superordinate) of objects was differentially related to the spatial scale of fMRI patterns, and that the spatial scale at which information was encoded increased along the ventral visual pathway. In sum, our results indicate that the proposed algorithm is particularly suited to assess and detect scale-specific information encoding in cortex, and promises further insight into the topography of cortical encoding in the human brain.

## Introduction

A major goal of human cognitive neuroimaging is to establish a mapping between mental representations and patterns of activity human cortex (van Essen et al. 2001; Logothetis & Wandell, 2004). The main description of this correspondence is functional localization, i.e. where on the two-dimensional cortical sheet neural representations reside (van Essen et al., 1998; Fischl et al. 1999; Brett et al., 2002). Neural representation in human cortex typically involves distributed neuronal populations. Thus, representations in neuroimaging are rarely restricted to single image points, but rather appear as patches of activation across the cortical sheet. Therefore, two further parameters of neural representations on the cortical beyond point location must be given: the spatial *scale* and the *form* of the pattern in the localized patch. Without information about spatial scale it remains impossible to correctly ascribe cognitive function to any of the multiple scales on which the brain is organized, ranging from single cells over cortical columns, patches and large-scale maps (Op de Beeck, 2008; Swisher et al., 2010; Brants et al., 2011; Misaki et al., 2013). Without a detailed characterization of the activation pattern, e.g. through the direction of a gradient, valuable and distinctive fine-grained information might be neglected (Portilla & Simoncelli, 2000).

The methodological challenge in characterizing the spatial patterns of human brain activity is that analysis must observe the structure restriction of a highly convoluted cortical sheet, and be carried out with respect to the underlying differential geometry of the irregular two-dimensional cortical sheet (van Essen et al. 2007; Chen et al. 2011), rather than three-dimensional Euclidean space (Brants et al., 2011). For this, two key technical challenges need to be addressed: 1) how to assess spatial scale on an irregular mesh that captures the geometry of the cortical sheet (Hagler et al., 2006) correctly, and 2) how to assess the directional components in the activation pattern (Simoncelli & Freeman, 1995).

Here, we address both issues simultaneously with an algorithmic scheme for directional spatial filtering on the cortical sheet. We built steerable bandpass filters on the irregular cortical surface, constructing differential operators of directional derivatives, and combining them with Gaussian smoothing kernels. To achieve an infinite-impulse response filter (IIRF) for Gaussian smoothing, we adopted a geometrical discretization of the Laplace-Beltrami operator (Meyer et al., 2003), combined with a modified algorithm for computing the symmetric matrix exponential (Sidje, 1998). Importantly, we note that the effective scales of information differ from the nominal filter sizes applied to extract it, due to the underlying smoothness of the data. Thus, filtering approaches must take this into account, and only the effective scales of information can be compared across different approaches.

We demonstrate the utility of the algorithm in comparison to previously proposed methods in two ways. First, using modelling we show that through improvement in the smoothing operations our proposed method yields superior results in numerical precision and spatial uniformity of filter kernels compared to the most widely adopted approach for cortical smoothing. Second, we apply the proposed method to an fMRI dataset to assess the cortical encoding of information about visual objects at the subordinate (exemplar) and superordinate (category) level and made several observations. We found that filtering by our method improved the detection of discriminant information about experimental conditions. Further, it provided a novel quantitative description of the spatial organization of encoding of visual categories: Information about ordinate level visual categories (e.g. distinguishing plane from car) was more prominent at a coarser scale than for subordinate categories (or exemplars, i.e. distinguishing one plane from another), and we observed a systematic increase in the spatial scale at which information was maximally explicit along the hierarchy of the ventral visual stream.

Together, this indicates that the proposed implementation to be particularly suited to assess and detect scale specific information encoding on the cortical surface, promising further insight into the topography of cortical encoding in the human brain.

## 1 Methods

### 1.1 Heat diffusion and Gaussian smoothing

Assessment of scale specific information relies crucially on the spatial smoothing operator and its implementation on the cortical surface. The smoothing operator must observe the geometry of the irregular mesh, and avoid introducing geometric distortions and inhomogeneity to allow for appropriate and unbiased assessment. Towards this aim we employed a Gaussian smoothing operator based on heat diffusion on irregular mesh.

#### 1.1.1 The relation of Gaussian smoothing to the heat diffusion equation

The Gaussian smoothing operation in space is mathematically equivalent to a temporal physical process of heat diffusion with the input signal as initial condition (Koenderink, 1984). The following partial differential equation characterizes this physical process:

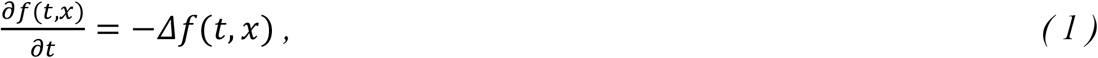

where Δ is the spatial Laplacian, or Laplace-Beltrami operator in case the diffusion process is on a differentiable manifold. The general solution to this equation, with initial condition *f*(0, *x*), can be given by:

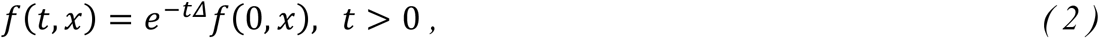

where *e*^−*t*Δ^, the diffusion operator, is the exponential of differential operator −*t*Δ. From the viewpoint of spatial smoothing filter, it is convenient to write above solution as:

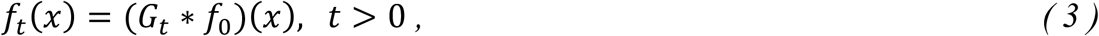

where *f*(*t*,·) = *f_t_* and *G_t_* = *e*^−*t*Δ^*δ*(*x*), the application of *e*^−*t*Δ^ to a Dirac delta function. The impulse function *G_t_* is also called the heat kernel, and the time variable *t* acts as the size or scale parameter of Gaussian smoothing kernel exp (−*x*^2^/*t*).

#### 1.1.2 Discretization of geometrical Laplace-Beltrami operator on triangulated mesh

When applied to a discrete surface mesh, the Laplace-Beltrami operator Δ needs to be discretized and expressed in matrix form. One of the most commonly adopted discretization of this differential operator is the so-called geometrical Laplacian: it takes the embedding geometry of the mesh into account and is given by (Meyer et al., 2003; Reuter, 2009; see Fig. 1A for a visualization of the parameters in the equations):

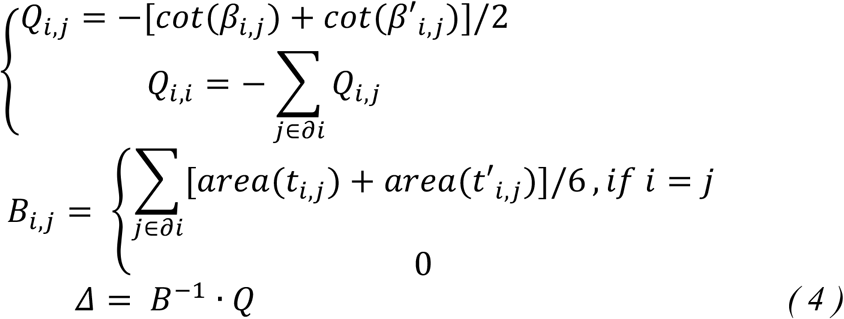

where *β* and *β*′ are the angles subtended by each edge, *t* and *t*′ the triangles at the two sides of each edge, and *∂_i_* indicates the immediate neighbors of vertex *i*. In practice, the Laplacian is implemented as a sparse matrix in which non-zero items correspond to edges in the mesh and are given as in Fig. 1A. Intuitively, each row of the symmetric matrix *Q* quantifies the conductivity relations between a vertex and its immediate neighboring vertices, whereas the diagonal matrix *B*, also called *lumped mass* matrix, specifies for each vertex a capacity factor, an integral measure for the vertex, so that the inner product of two functions on the underlying surface 〈*f_x_, f_y_*〉 = ∫_*M*_ *f_x_* · *f_y_ds* can be numerically computed by 〈*x, y*〉 = *x^T^By*.

**Figure 1:**
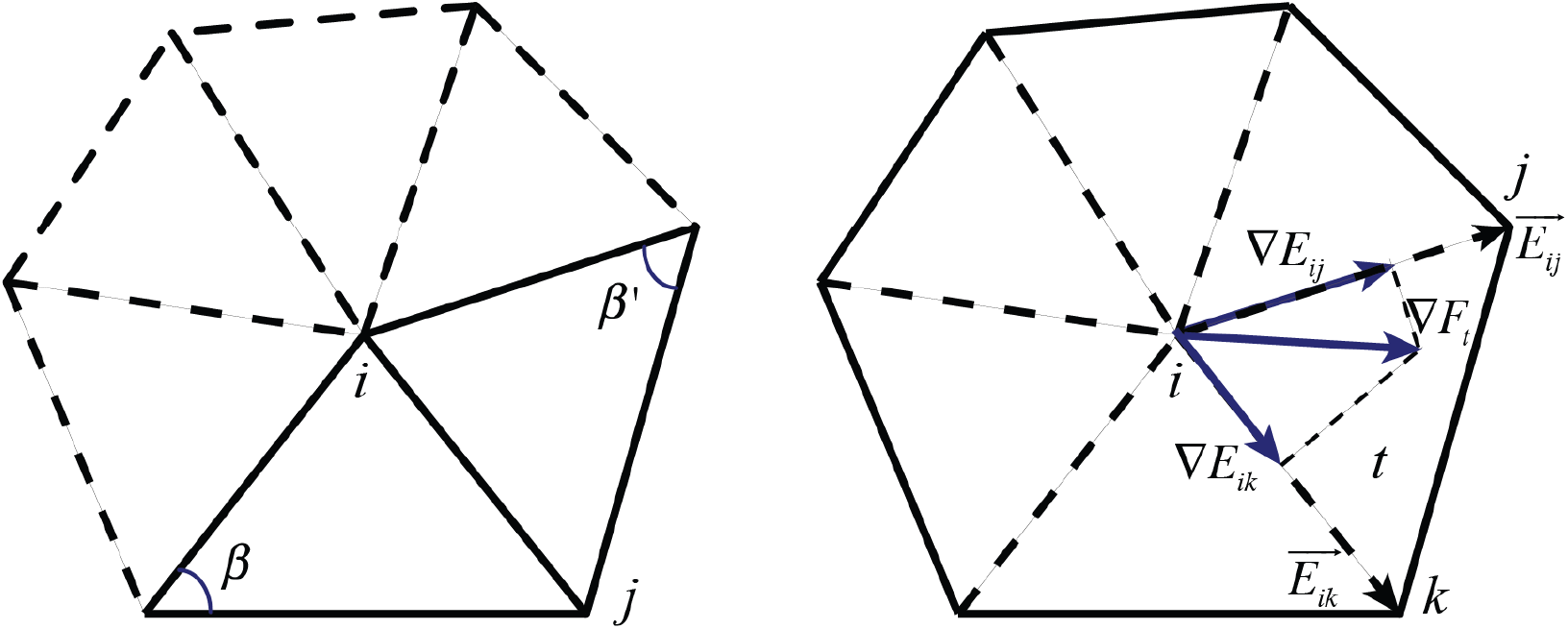
Geometric Laplacian and directional gradient on surface mesh. (A): the parameters of the discrete Laplacian-Beltrami operator on a triangulated mesh for the *i*-th vertex, as in (4). (B): parameters for estimation of gradients for defining directional derivative operators as in (9).

#### 1.1.3 Calculating numerical solutions to the diffusion equation

For a given input function *f*_0_ and a scale parameter *t*, we can substitute (4) into (2) and use a matrix exponential algorithm to compute the numerical solution *f_t_* by:

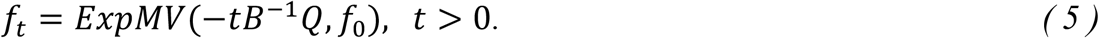

where *ExpMV*(*A, ν*) approximates exp (*A*) · *ν* without computing exp (*A*) explicitly (Sidje, 1998). See **Supplementary Text** for more detail about this algorithm and an efficient implementation for diagonal *B* and symmetric *Q*.

#### 1.1.4 Laplacian of Gaussian as bandpass filters

As from (3), the solution *f_t_* approximates the smoothing of input *f*_0_ by a Gaussian kernel of size *t*. Applying the Laplacian *B*^−1^*Q* to *f_t_*, we can immediately get the bandpass filtered detail of *f*_0_ at scale of parameter *t*, with respect to the symmetric, second derivative of Gaussian:

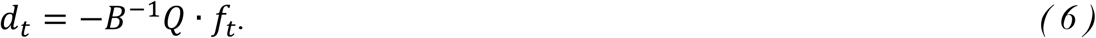

We note (6) is the ubiquitous feature detector in computer vision algorithms (Marr and Hildreth, 1980), Laplacian of Gaussian (LoG), in form of a discrete differential operator on discrete surface. Notice the equivalence of the right sides of (6) and (1): From the perspective of scale space representation, LoG simply acts as the partial derivative of a multiscale function with respect to its scale parameter.

### 1.2 Steerable filters of directional derivatives of Gaussian

#### 1.2.1 Local directions are necessary for defining directional derivative operators

To construct steerable bandpass filters at specific scales, we first note the differential property of convolution:

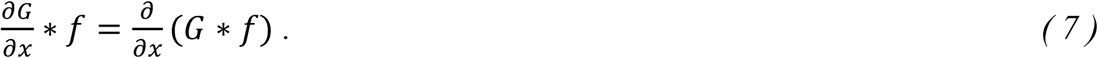

Therefore, if we have already computed *f_t_* by applying a Gaussian kernel *G_t_* to an input function *f*_0_, we can simply apply a differential operator to *f_t_* to get the scale-specific details of *f*_0_, equivalent to the outputs from bandpass filters of Gaussian derivatives. Particularly, as we are concerned with functions defined on a 2D manifold, we would like to have differential operators for partial derivatives in orthogonal directions on the surface, so that the linear combinations of them could be “steered” to any possible direction in the tangent bundle of the surface. This property of orthogonal directional derivatives is called steerability (Freeman and Adelson, 1991; Simoncelli & Freeman, 1995)

To construct such differential operators for directional derivatives, we need to define a system of directions at every vertex on the surface mesh. These directions should be uniformly consistent: The directions over neighboring vertices being parallel to each other. Geometrically, this is equivalent to planar parameterization of the surface, and is only possible for surfaces with zero Gaussian curvature everywhere. For our application, however, it may suffice to define such directions that are parallel to each other over flat area and change smoothly and consistently over a curved area.

#### 1.2.2 Gradients of Fiedler vector field as local directions

We choose to define these directions by using the discrete gradients of the Fiedler vector *F*_Δ_ (Biyikoglu et al. 2007), defined as the generalized eigenvector corresponding to the 2nd smallest eigenvalue *λ* of the discrete Laplace-Beltrami operator Δ:

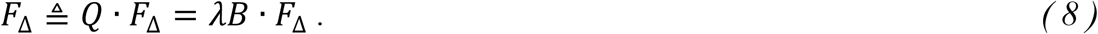

When the underlying surface is sufficiently smooth, the Fiedler vector is the smoothest bi-modal function defined on the vertices and its gradient field ∇*F*_Δ_ is consistent almost everywhere (except at very few modal and saddle vertices).

#### 1.2.3 Approximation of Fiedler vector gradients on mesh and directional derivative operators

To calculate the gradient of Fiedler vector *F* at the vertices on a triangulated mesh, we assume piece-wise linearity of the underlying Fiedler function on the triangle faces so that the gradient on a triangle is constant and can be computed by linear fitting:

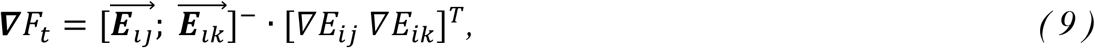

where *i,j,k* are the vertices of the triangle face *t*, 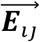 and 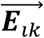 are the normalized edge vectors, ∇*E_ij_*, ∇*E_ik_* the gradients of *F* along the two edges and []^−^ the pseudo inversion of matrix (See Fig. 1B for the parameters in the equation).

Note that the above procedure for calculating the gradient of *F* is applicable to any smooth function *f* defined on the surface, thus on each triangle face, the partial derivatives of a given function *f* along the direction of the gradient of Fiedler vector can be calculated via the inner product of the two gradients:

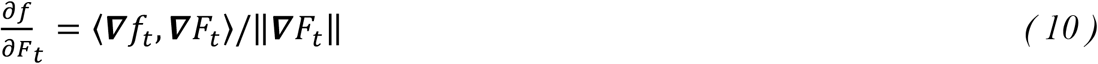

The directional derivative of *f* at vertex *i* is then estimated by the area-weighted average of the partial derivatives on all the triangles containing vertex *i*:

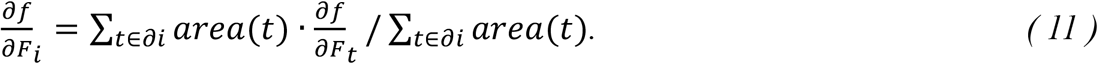

Furthermore, by defining the orthogonal direction of the Fiedler gradients as the cross products of them and the face normal vectors:

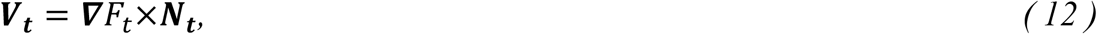

where ***N_t_*** is the normal vector of the triangle face, we can compute the directional derivatives on this orthogonal direction in the same way as on the direction of Fiedler gradients. It is important to note that these two orthogonal directions (hereafter referred as primary and secondary directions) thus allow us to construct directional derivative operator for each vertex, on any local direction in the plane tangent to the vertex, by simply taking a proper linear combination of them.

Fig. 2A shows the Fiedler vector on a cortical surface mesh with a zoomed-in portion showing the locally defined primary directions. In Fig. 2B, filters of directional derivatives of Gaussian are visualized with their impulse response functions.

**Figure 2.**
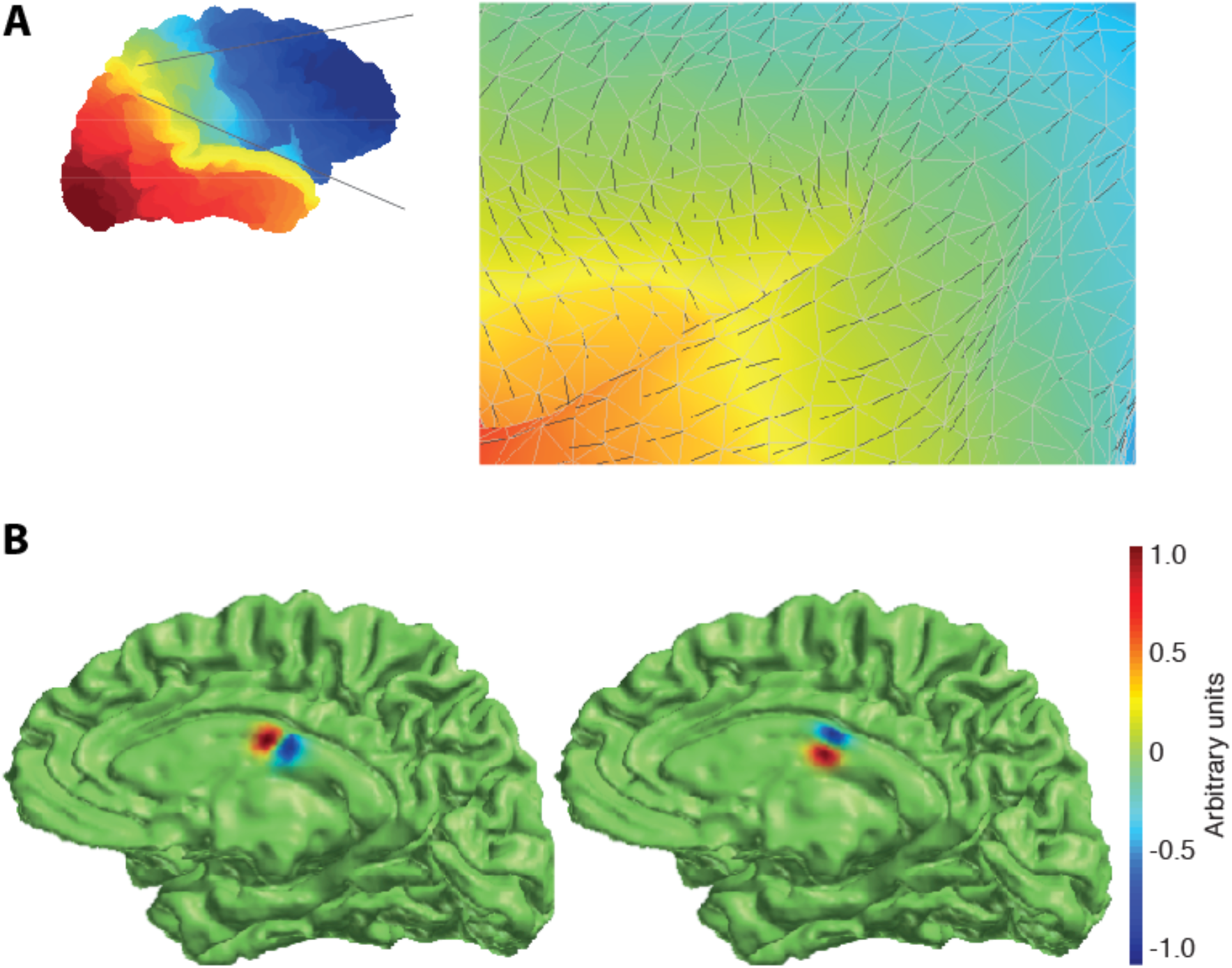
Directional derivatives of Gaussian based on directions defined by Fiedler vector. (**A**) Visualization of Fiedler vector of the discrete Laplacian-Beltrami operator on a patch of cortex (indicted by black arrows). Colors indicate position in space along the posterior-anterior direction. (**B**) Impulse responses of the filters based on directional derivatives of Gaussian, normalized to unit numerical range (left: primary direction; right: secondary direction). Colors in arbitrary units indicate filter weights.

### 1.3 Effective filter size and effective scale

#### 1.3.1 Effective filter size is estimated from the smoothness of its action on Gaussian random field

In order to build a pyramidal representation with linearly growing spatial scale, we need to determine the scale parameters *t* of the heat diffusion kernel to relate its value to smoothing filter size. Here we followed the practice in Hagler et al. (2006) by estimating the overall smoothness of filtered Gaussian random noises as the equivalent full-width-at-half-magnitude (FWHM) size of these filters. Specifically, we generated independent, uniformly distributed random noise on the surface, applied the filters to it and estimated the smoothness according to random field theory (RFT):

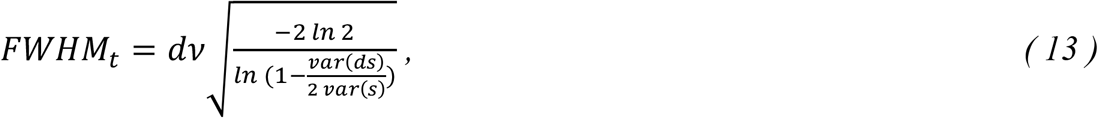

where *dv* is the average edge length, var(*ds*) the variance of difference between neighboring vertices, and var(*s*) the total variance over all the vertices. Note the FWHM for Gaussian smoothing kernel exp (−*x*^2^/*t*), is proportional to the square root of the scale parameter *t*. Therefore, we calculated the FWHM for each cortical surface mesh on a range of parameters *t*, and took the linear fitting of it and 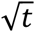 to extrapolate for other filter size regarding parameter *t*. The FWHM calculated in this way is taken as the *effective filter size*.

#### 1.3.2 Effective scale is estimated from the smoothness of residual data

While the effective filter size can be a valid estimation of spatial scale for functions that are smoothed from independent Gaussian random noise, the surface images mapped from volume data may often violate the independence assumption. To estimate the *effective scales* of the results, we opted to adopt a *post hoc* estimation, by calculating the ratio between the cortical surface area and the number of resels computed by SurfStat from the *residuals*:

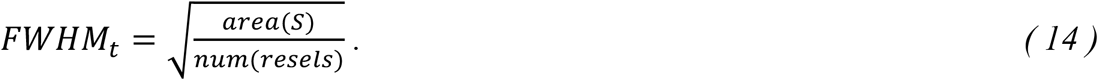

Note the resels returned from SurfStat are multi-dimensional and only that of 2D, or areal resel number, is used in (14).

### 1.4 Evaluation of Gaussian smoothing algorithms

In order to evaluate the numerical precision and spatial uniformity of the proposed heat diffusion smoothing algorithm, we applied it to impulse functions on a sphere mesh, on which Gaussian kernels can be calculated analytically and then sampled for reference. We created sphere meshes by iteratively subdividing a regular tetrahedron and projecting new vertices to the sphere. In doing so, we constructed a topologically almost-everywhere regular mesh: All except the initial 4 vertices have the same connectivity of 6. On the other hand, geometric irregularity of variable areal measures is introduced by the spherical projection. An elastic regularization was applied in each of the iteration to control this areal variability. We repeated this iterative procedure for 7 times to generate a sphere mesh of about 32,000 vertices (radius: 10 mm, average edge length: 0.1385 mm). Impulse functions at random locations on the sphere are then generated and filtered by different smoothing algorithms for comparison.

### 1.5 FMRI experiment and data preprocessing

To demonstrate the approach used here we re-analysed data from an fMRI experiment on categorical-level and exemplar-level representation of visual objects (published previously in Cichy et al., 2011). We only give a briefly summary here. 13 healthy subjects (1 subject’s data were not included in this analysis due to poor T1/EPI volume alignment) participated in a mini-block (duration: 6s) design. Stimuli were 3 different exemplars from 4 different categories (animal, chair, car and airplane), yielding a total of 12 different images. In each mini-block, a single object was rendered in 3D (6 renderings presented for 800 ms with 200 ms gap) at a position either 4° right or left of the screen center, subtending ~4.6° of visual angle. Each rendering either repeated the previous viewpoint, or displayed with a random viewpoint at least 30° difference in rotation in depth compared to the previous rendering. The number of repetitions of viewpoints was counterbalanced across objects. Subjects were instructed to fixate at the center of the screen and perform a one-back viewpoint judgment task.

Functional images were acquired with a gradient-echo EPI sequence (TR = 2000 ms, TE = 30 ms, flip angle = 70°, FOV = 256 mm, matrix = 128 × 96, interleaved acquisition, no gap, 2mm isotropic voxels, 24 slices). Slices were positioned along the slope of the temporal lobe to cover the ventral visual cortex. Each run of the main experiment has 412 volumes; in total 5 experiment runs were collected for each subject. In addition, a whole brain EPI volume was also acquired in a separate run to facilitate the T1/EPI alignment. All functional volumes were motion corrected using SPM8, and aligned to the whole brain EPI volume, which was coregistered to the structural volume. Realignment parameters were later used in hemodynamic modeling to eliminate motion-induced artifacts.

### 1.6 Cortical surface mesh generation and volume-surface data mapping

Cortical surface meshes were generated for each subject from high-resolution structural MRI scans (192 sagittal slices, TR = 1900 ms, TE = 2.52 ms, flip angle = 9°, FOV = 256 mm, 1 mm isotropic voxels) with FreeSurfer version 5.1 (Dale et al., 1999; Fischl et al., 1999). A gray-mid layer lying half the distance between white matter surface and pial surface was created for volume-surface data mapping, as it has optimal uniformity of surface curvature and offers good balance between spatial specificity and sensitivity of information extraction (Chen et al. 2011). To avoid oversampling in data mapping, we further simplified the generated mesh using CGAL library (www.cgal.org), to make sure that all the length of the mesh edges are between 1 and 2mm. This simplification also reduced the number of vertices up to 50% and speeded subsequent analyses remarkably. The raw volume data were then tri-linearly sampled with the vertex coordinates to complete the volume-surface mapping, so for each volume we had a discrete function defined on the vertices, which is called *surface image* hereafter.

### 1.7 Multivariate statistical analysis of discriminant information

#### 1.7.1 Temporal and spatial filtering on surface images

For each vertex, the values from all the surface images constitute a time series. We first applied temporal highpass filtering and pre-whitening to these time series, vertex-by-vertex, using SPM8. Heat diffusion smoothing was then applied to the surface images, time point by time point, with pre-computed scale parameters. At each scale and to each smoothed surface image, differential operators of directional derivatives and symmetric Laplacian were applied to extract the scale-specific details. This procedure made available for us both the pyramidal representation and the scale-specific details. Note that while the outputs from smoothing and symmetric Laplacian of Gaussian filtering are univariate, the outputs from the directional derivative filtering are bivariate.

#### 1.7.2 GLM estimation

Next, we modeled the cortical response to the 24 experimental conditions (12 objects presented either in the left or the right hemifield). To estimate the overall smoothness of residuals, all the five runs in the experiment were modeled together for each subject. The onsets of the mini-blocks were entered into the general linear model (GLM) as regressors of interest and convolved with a canonical hemodynamic response function (HRF). All these regressors of interest, together with that of the motion parameters and default baseline, were also preprocessed with temporal highpass filtering and pre-whitening. We then fitted the preprocessed GLM to the spatially filtered data, at each scale and vertex by vertex, to estimate the model parameters and residuals, which were later used for calculating the effective scales.

#### 1.7.3 Using SurfStat for multivariate analysis and smoothness estimation

To investigate the scale-specific information that differentiates the categories of objects, particularly for the bivariate details extracted by the directional derivative filters, we used the SurfStat toolbox (Worseley et al., 2009) to compute the *F*-statistics on two categorical levels: On the subordinate level, the null hypothesis assumes that all the 3 exemplar objects within the same category have the same mean over runs; at the ordinate level, the null hypothesis assumes that all the 4 categories have the same mean, where the 3 objects within each category were treated as repeated observations. In both cases, we treat the presentations in different hemifields as repeated observations of the same object. Note for multivariate parameters, SurfStat computes the Roy’s greatest root as the *F*-statistic, which is the largest *F*-value over all possible linear combinations of the input variables. The statistical significance of the results and the respective significance thresholds regarding surface-based multiple comparison correction, is also derived by the routines in SurfStat.

## 2 Results

### 2.1 Comparison of smoothing quality by heat diffusion smoothing versus smoothing through iterative averaging

To evaluate the quality of the heat diffusion smoothing operator, we applied it in a model case for which analytic solutions are readily available, and compared the results to the smoothing operator based on iterative averaging, i.e. the current standard procedure as implemented in Freesurfer.

In detail, we generated 100 impulse functions at random locations on a sphere mesh. For each location, a Gaussian kernel with unit sigma was calculated and sampled to the vertices as a *discretized Gaussian* for reference. We then applied the heat diffusion and the iterative averaging algorithms to the impulse functions and calculated the mean squared errors (MSE) with respect to the discretized Gaussians. Note that for comparison across smoothing approaches, the smoothing parameters, i.e. the effective filter sizes, have to be the same, which were determined with RFT-based estimation of smoothness before the comparison.

We made two observations. First, we found that the MSE between heat diffusion smoothing and the reference discretized Gaussian in both absolute and relative terms was ~30 times smaller than iterative averaging (Table 1A). Fig. 3 displays representative results of iterative and heat diffusion smoothing, demonstrating this point visually. Second, we observed that for heat diffusion smoothing the variance of filter sizes was comparable to the reference discretized Gaussian, while it was ~10 times larger for iterative averaging (Table 1B). In Figure 3, this is expressed visually by the fact that the smoothing results from heat diffusion converge not only more geometrically to the discretized Gaussians, but also more uniformly over regions of different triangulation density. In contrast, the iterative averaging introduced remarkable geometric distortion and inhomogeneity.

**Figure 3:**
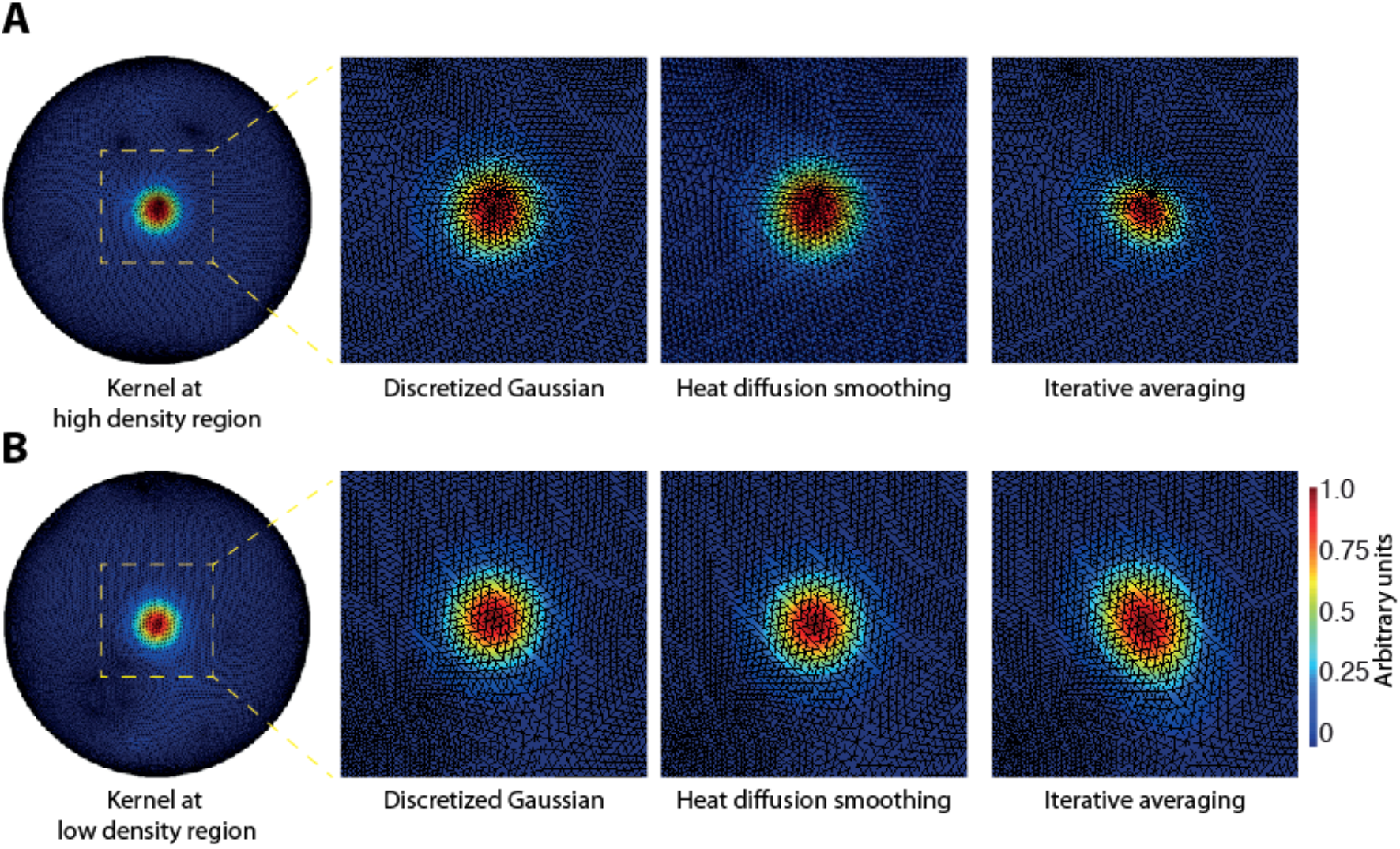
Gaussian smoothing on a sphere mesh at regions with different triangulation density. Left: Discretized Gaussian; Middle: Heat diffusion smoothing; Right: Iterative averaging. Results from an input of impulse function located at regions of high (A) and low (B) density of triangulation. Compared to heat diffusion smoothing, iterative averaging introduces density-dependent size inhomogeneity and geometric deviation from discretized Gaussian.

**Table 1:** Approximating precision of smoothing algorithms with respect to (A) error and (B) filter size. Filter parameters were first determined by matching the RFT smoothness to the FWHM of sampled Gaussian. Mean squared errors are averages over 100 smoothing results regarding the respective sampled Gaussian. Filter sizes were then empirically estimated by the square root of the area of vertices with value greater than half of the maximum, the variance of size is calculated over 100 instances.

|  | (A) |  | (B) |  |
| --- | --- | --- | --- | --- |
|  | Abs. Error | Rel. Error | Filter Size | Var. of Size |
| Sampled Gaussian |  |  | 2.1089 | 0.0357 |
| Heat Diffusion Smoothing | 0.0003 | 0.0054 | 2.0980 | 0.0411 |
| Iterative Averaging | 0.0109 | 0.1816 | 2.1321 | 0.3257 |

Together, our results show that heat diffusion smoothing provides higher numerical precision and geometric uniformity than iterative averaging. Thus, for further filtering analyses on the cortical surface we used only heat diffusion smoothing.

### 2.2 Comparison of different filtering operations on the cortical surface in revealing discriminative information

To assess the spatial scale at which information is encoded on the cortical sheet, activation patterns must be filtered at different spatial scales. Here, we evaluated three types of smoothing filters. First, we used Gaussian smoothing (SM) in the heat diffusion implementation at different scales, resulting in low-pass filtered activation patterns. Second, to isolate a specific spatial scale beyond simple low-pass filtering, we used Laplacian of Gaussians (LoG) as a band-pass filter. The result of LoG filtering are band-passed activation patterns. Third, to also take into account that spatial patterns on the cortical surface have gradients and orientations, we used directional derivatives of Gaussians (dDG). The result of dDG filtering are band-passed and direction-specific activation patterns.

#### 2.2.1 Matching effective filter size is a crucial precondition for comparing results of filtering approaches on the cortical surface

A precondition for a proper comparison of the results of the proposed filtering methods is that results are compared when the same filter sizes are compared. However, as the effective size is estimated by the pattern smoothness with RFT-theory (see 1.3), *effective* filter sizes might differ from *nominal* filter sizes when the elements of the smoothed patterns are spatially correlated. More specifically, spatially correlated patterns would decrease the variance of neighbouring difference *ds* in (13), thus increase the overall estimation. As fMRI voxels that make up activation patterns do show strong dependence, it cannot be assumed that effective and nominal filter sizes are identical. To determine the relation between the diffusion parameter and the resultant effective filter size on cortical surfaces, we generated 100 normally distributed random functions on each surface, and applied heat diffusion smoothing with parameter *t* ranging from 2 to 36. Fig. 4A shows the effective filter size in relation to the square root of *t*, as estimated from RFT-based smoothness. We observe that, compared to the application of Gaussian smoothing operator, the application of differential operators of either geometric Laplacian or directional derivatives *decreases* the RFT-based smoothness estimation of the effective filter size.

**Figure 4:**
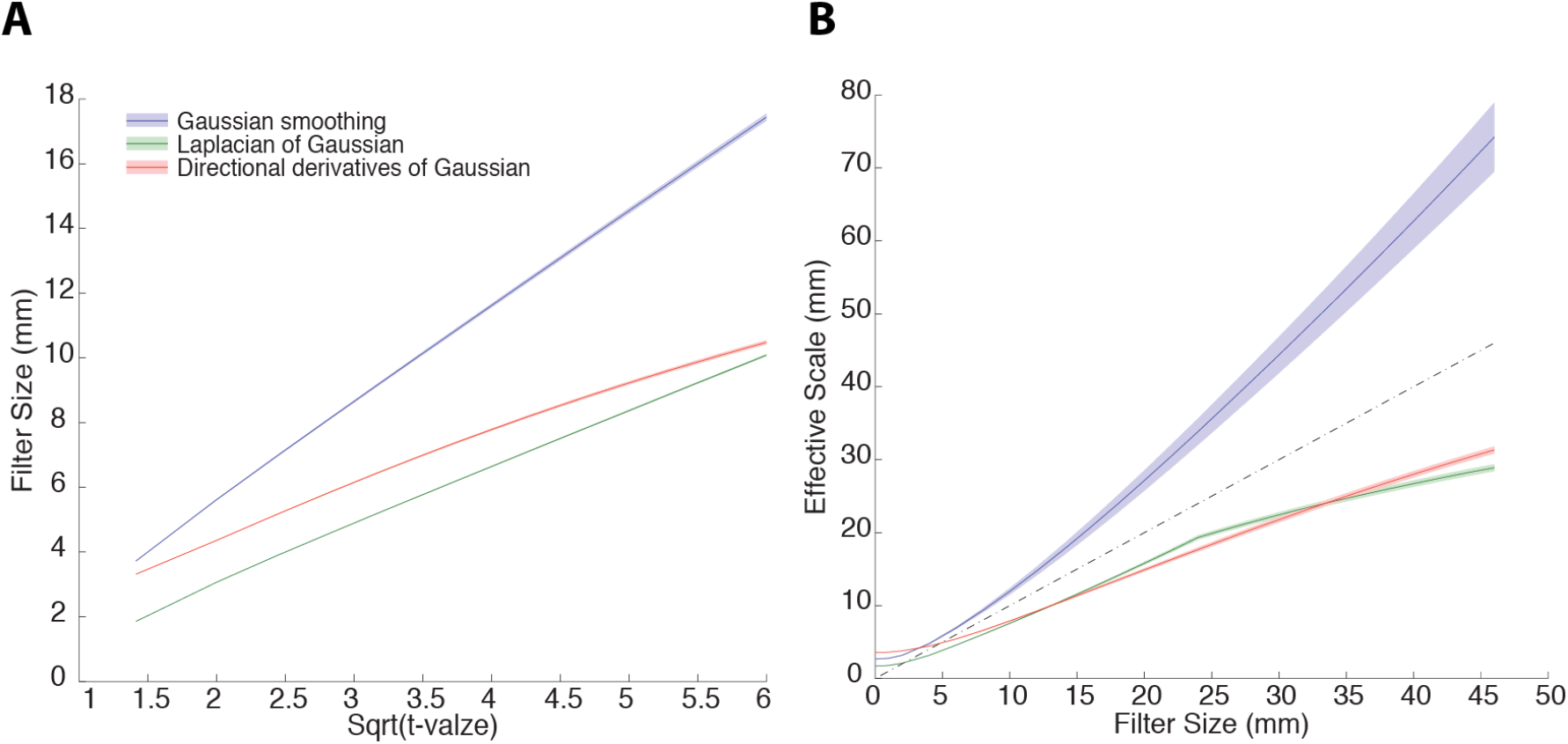
Effective filter size and effective scale. (a): Effective filter sizes as estimated from RFT smoothness, plotted against the square root of diffusion parameter *t*, for Gaussian smoothing (SM), Laplacian of Gaussian (LoG) and directional derivatives of Gaussian (dDG). Application of Laplacian of Gaussian or directional derivatives of Gaussian decreases the RFT-based smoothness estimation of the effective filter size, requiring correction. b): Effective scales of the residuals from the three different kinds of filtering of fMRI data on the cortical surface, plotted against the effective size of SM filter. The dash line shows the equality line of the effective scales and the effective filter sizes. In both plotting, the shaded area indicates the range of standard error across 12 subjects.

Thus, we equated effective filter sizes before comparing results from different filtering methods based on a post-hoc estimation of smoothness of fMRI data (Fig 4B). For the analysis of the spatial scale at which information is encoded on the cortical sheet, we used a linear range of effective filter sizes of SM from 0 to 46 mm (size 0 for no smoothing), in 2 mm steps. In Fig. 4B we plotted the effective scales estimated from the resel numbers of residuals computed by SurfStat toolbox, against the effective filter sizes of Gaussian smoothing (SM). Corroborating the results of modeling, we observed that the effective scales of residual were noticeably greater than the effective sizes of the smoothing filters (Hagler et al. 2006), but smaller than that of the differential operators being applied.

### 2.3 The distribution of information on the cortical sheet as resolved by different filtering operators

We compared the ability of SM, LoG and dDG to reveal the nature of fMRI activation patterns underlying information encoding in human visual cortex. For this, we used an fMRI data set mapping activity in ventral visual cortex while participants viewed 3 different object exemplars in 4 different categories (cars, chairs, planes and animals), i.e. in total 12 different objects presented to the left and right of fixation. This allowed us to determine the spatial scale at which information about objects is encoded at two levels of abstraction: the ordinate category level (e.g. car vs. plane) and the sub-ordinate level (e.g. one car vs. another car). To determine information encoding, we used multivariate pattern classification.

First, we assessed the spatial distribution of information about objects in a spatially unbiased analysis. That is, we determined discriminant information between objects on the cortical sheet detected by multivariate analysis for the three different filtering methods (SM, loG, dDG) at two levels of abstraction (sub-ordinate and ordinate category level). Representative results for a single subject at two different spatial scales (equalized effective scales) are plotted in Fig. 5.

**Figure 5:**
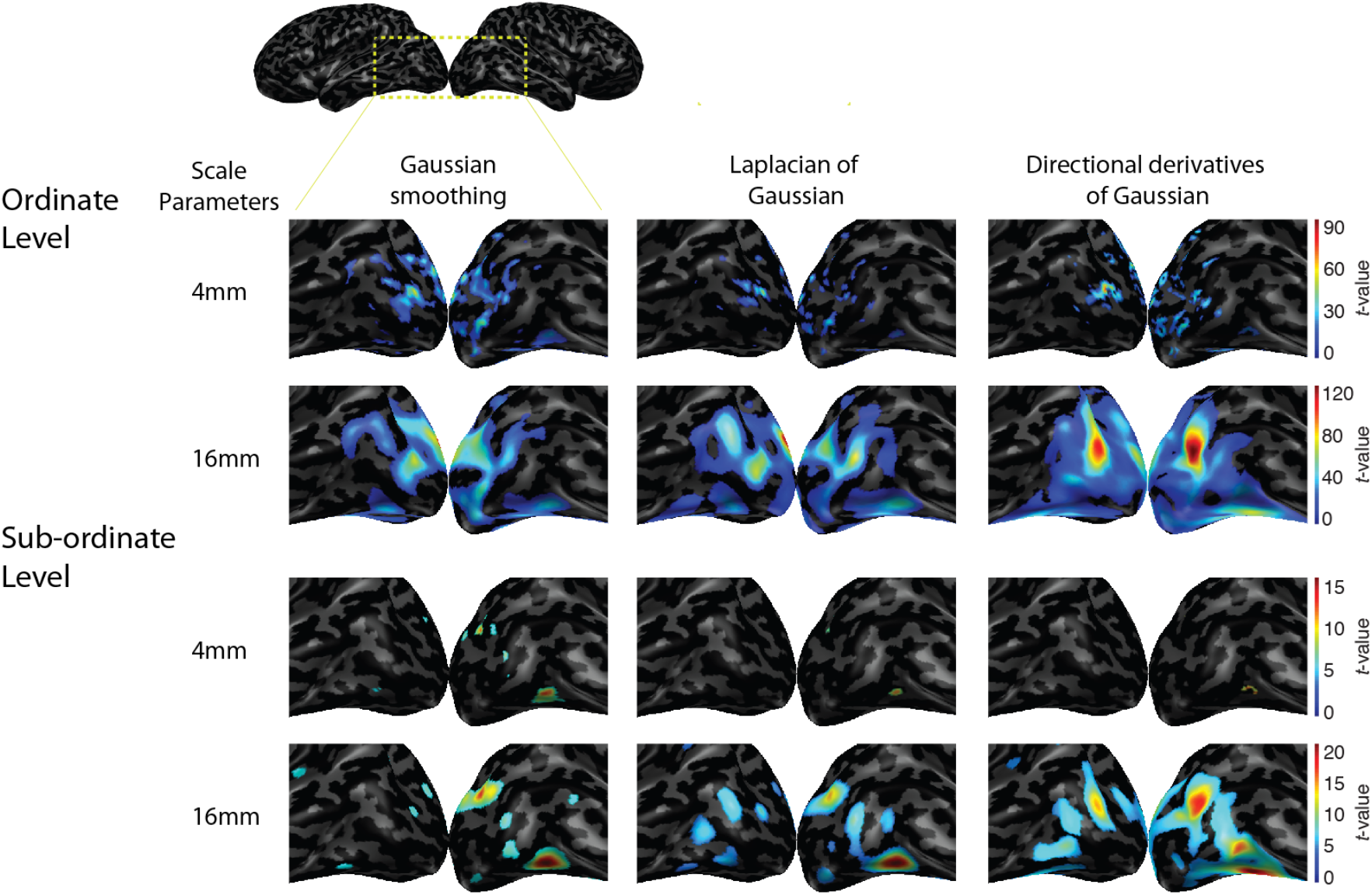
Map of discriminant information about ordinate and subordinate categories at two different scales (fine: 4mm; coarse: 16mm). F-statistics (thresholded at *P* < 0.01, FWE corrected) from one subject are rendered on inflated cortical surface and the lateral-occipital portion of the ventral visual cortex is highlighted by the zoomed-ins. For comparison, results from SM (left), LoG (middle) and dDG (right) are presented side by side, with colors normalized to the same range for each row. We make three qualitative observations: First, SM, LoG and dDG yield increasingly statistically significant results, suggesting the bandpass filters and directional filters outperform highpass and symmetric filters in revealing encoded information in cortex. Second, while coarser (16mm) filtering yields stronger results that finer (4mm) filtering, the relative difference depends on the level of categorization. Third, filtering at 4mm yields more posterior results than filtering at 16mm, suggesting that spatial scale at which objects are encoded in ventral visual cortex might increase from anterior to posterior.

For all filtering operations, the regions containing significant discriminant information about objects include occipito-temporal cortex on the lateral and ventral surface of the brain, in line with previous studies reporting the location of object representations (Cichy et al. 2011; Chen et al. 2011). However, we also note three qualitative differences between filtering operations: overall the results from LoG appear stronger (i.e., yield higher statistical values and effects of larger extent) than for SM, and stronger for dDG than for LoG, suggesting that bandpass filters outperform high-pass filters in revealing discriminant information, and so directional over symmetric filters. Second, while in general discriminative information seems to be higher for coarser filtering (16mm) compared to finer (4mm) filtering, the results of the filtering operations differ in the relative strength depending on whether information pertains to sub-ordinate and ordinate level. Thus, the filtering methods might be differentially sensitive in detecting differences in spatial scales at which discriminative information for ordinate vs. sub-ordinate category distinction is encoded in the brain. Third, results from filtering at 4mm appear more prominent in posterior portions of the visual brain compared to filtering at 16mm. This suggests that the spatial scale at which object information is encoded in ventral visual cortex might increase from posterior to anterior. For quantitative assessment across subjects, we investigated each of those three observations further in a region of interest analysis below.

#### 2.3.1 Bandpass filtering improves discriminant analysis power of multivariate fMRI analysis

We investigated quantitatively whether LoG, dDG and SM differ in the strength of discriminant effects across subjects in a region-of-interest (ROI) analysis. We defined ROIs anatomically based on Freesurfer parcellation covering the lateral and ventral surface of occipito-temporal cortex from the occipital pole to inferior temporal cortex (Fig. 6A). To assess possible posterior-to-anterior gradients in information encoding along the ventral visual pathway, we split three parcellations (lingual, lateral-occipital and fusiform gyrus) into anterior and posterior parts. This resulted in 8 ROIs in total, ordered in posterior-to-anterior direction: pericalcatrine cortex (PC), anterior and posterior lingual cortex (aLN, pLN), anterior and posterior lateral-occipital cortex (aLO and pLO), anterior and posterior fusiform cortex (aFF, pFF), and inferior temporal cortex (IT).

**Figure 6:**
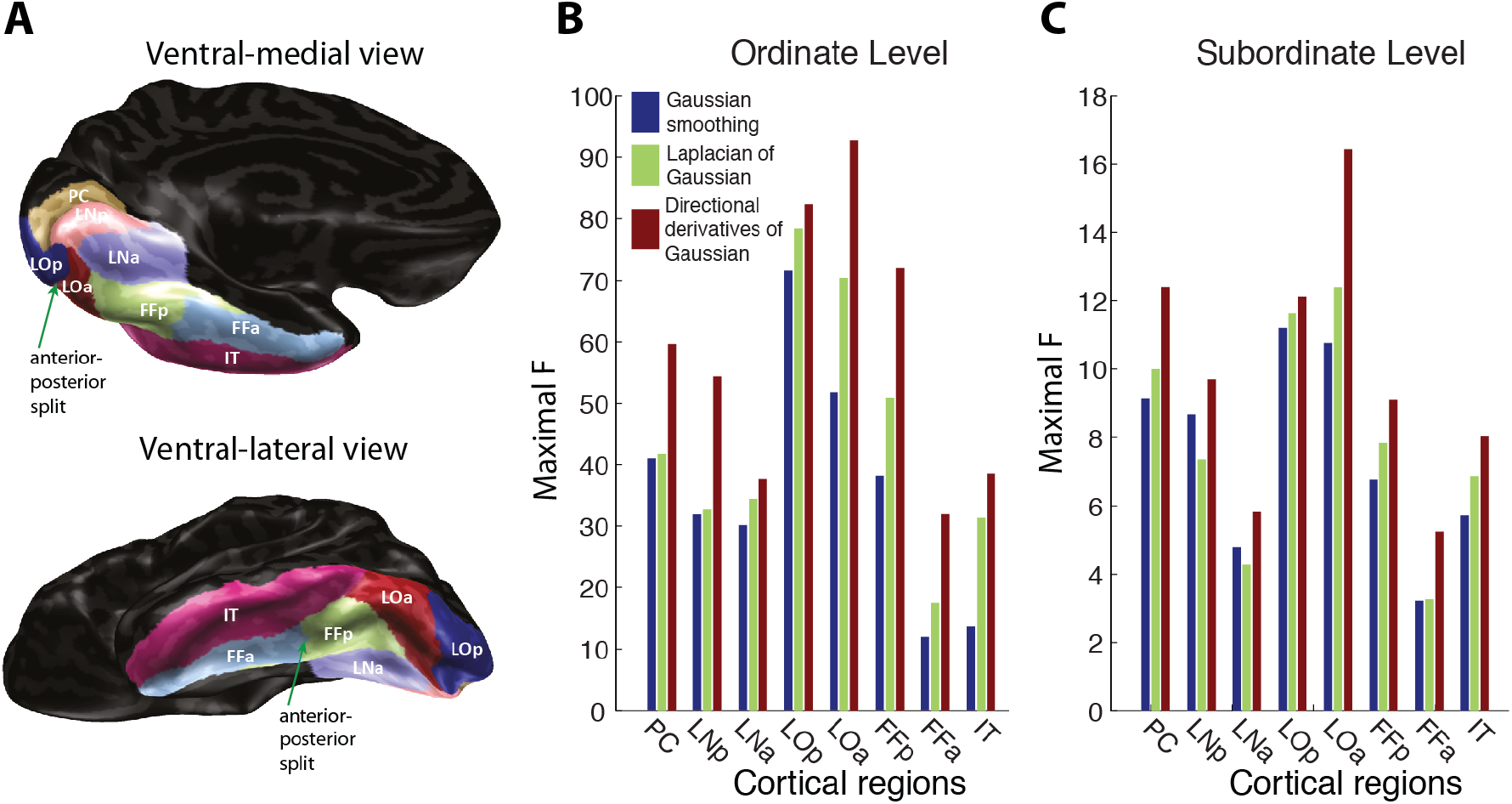
Regional maximal F-statistics. Maximal *F*-statistics across scales are shown here for discriminant information on ordinate (left) and subordinate categories. Medians instead of means across subjects are plotted, due to the fact that *F*-statistics are ratios of Chi-square statistics and not subject to direct summation. **Inset**: Anatomical regions for the analysis. Medial (left) and lateral (right) view of the ventral visual area anatomically parcellated by Freesurfer. Three regions were further split into anterior and posterior part as indicated by the green arrows. In total 8 regions were used in subsequent regional analyses: Pericalcarine cortex (PC), anterior and posterior lingual cortex (LNa, LNp), anterior and posterior lateral-occipital cortex (LOa, LOp), anterior and posterior fusiform cortex (FFa, FFp) and inferior temporal cortex (IT) (**B**) Group results (mean scales over subjects) from three different filtering methods: Gaussian smoothing (SM, left); Laplacian of Gaussian (LoG, middle) and directional derivatives of Gaussian (dDG, right). Corroborating the qualitative observation, we found significantly higher F-statistics from bandpass filtering (LoG, dDG) over smoothing (SM) in many ROIs, and for directional (dDG) over symmetric (Log) filtering (for details see Table 2).

Figure 6 shows the maximal *F*-statistics for object discrimination at the ordinate (Fig. 6b) and the subordinate (Fig. 6c) level across subjects for each filtering operation for each ROI. Concurrent with the qualitative observation from information maps as reported in Fig. 5, we found significantly higher F-statistics (Wilcoxon signed-rank tests) from bandpass filtering (LoG, dDG) over smoothing (SM) in many ROIs, and for directional (dDG) over symmetric (Log) filtering (for details see Table 2). Together, these results demonstrate the increased power of bandpass filters over simple smoothing to reveal discriminant information in spatial activation pattern on the cortical sheet. Please note that these peak *F*-statistics are maximal over all the scales, implying that bandpass filtering as a discriminant information detector can outperform *any* size of smoothing. Further, our results highlight the additional value of assessing the direction of gradients in activation patterns for increased discrimination performance.

**Table 2:**
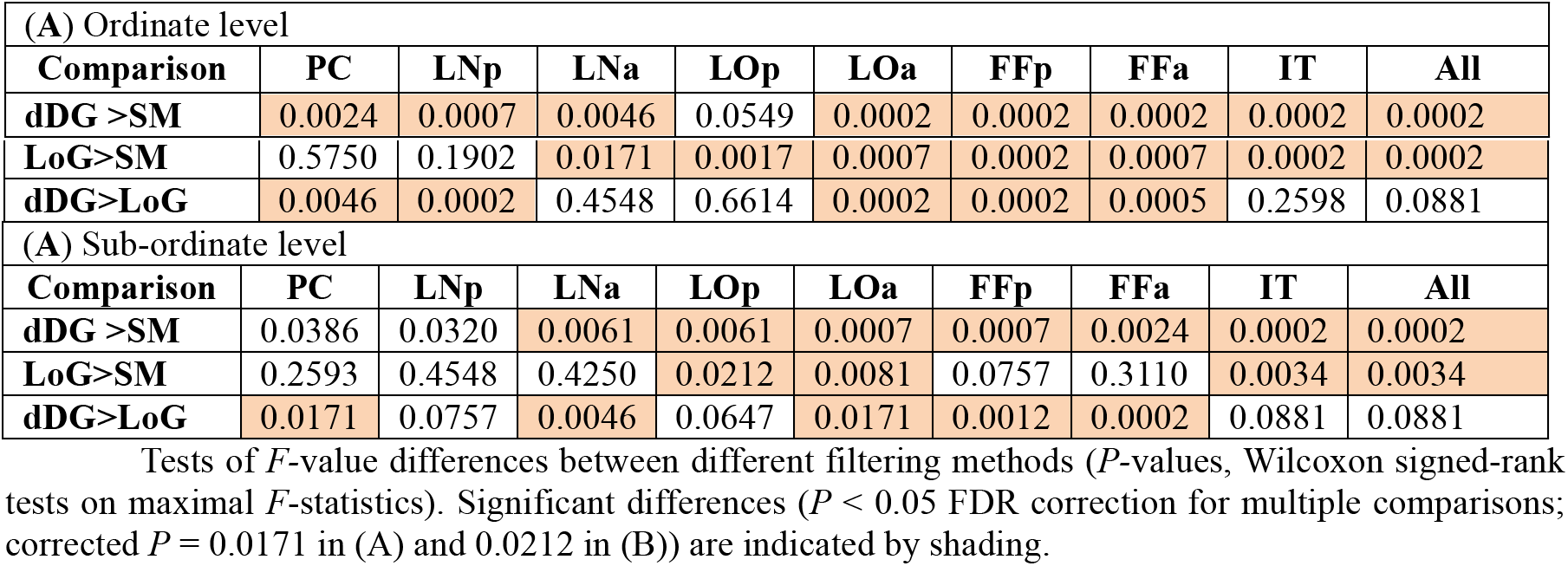
Tests of *F*-value differences between different filtering methods (*P-*values, Wilcoxon signed-rank tests on maximal F-statistics). Significant differences (*P* < 0.05 FDR correction for multiple comparisons; corrected *P* = 0.0171 in (A) and 0.0212 in (B)) are indicated by shading.

#### 2.3.2 Only directional derivatives of Gaussian uncovers differences in spatial scale of cortical information encoding for different levels of abstraction

Qualitative inspection of the information maps suggested that filtering methods might be differentially sensitive in detecting differences in spatial scales at which discriminative information for the ordinate vs. sub-ordinate category distinction is encoded in the brain.

Here we further investigated this observation quantitatively, assessing the propensity of high-pass smoothing (SM), Laplacian of Gaussian (LoG) and directional derivatives of Gaussian (dDG) to reveal those differences. For this we determined the effective scales for which the *F*-value distinguishing conditions at the sub-ordinate or ordinate level was maximal for each ROI and each filtering method (Fig. 7a for SM, 7b for LoG and 7c for dDG). To evaluate significance of differences in the spatial scale at which information is encoded at the sub-ordinate vs. the ordinate level, we conducted a 3×8 two-way ANOVA with factors filtering method (SM, DDG, LoG) and ROI (PC, pLN, aPL, pLO, aLO, pFF, aFF, IT). We found that the main effect for method was significant (*F* = 7.70, *P* = 0.0006), but not the main effect of ROI (*F* = 1.65, *P* = 0.1215), and the there was no interaction (*F* = 1.62, p = 0.0749) between the method and the region factors. We thus collapsed data across ROIs, and tested for differences in effective scale by method in two-sample t-tests (FDR corrected for multiple comparisons). We found that only dDG (*P* = 0.001), but neither SM (*P* = 0.99) nor LoG (*P* = 0.40) revealed a significant difference between categorical levels. Further direct comparison of filtering methods by paired t-tests (*P* < 0.05, FDR corrected) revealed an order with respect to the differences in resolving spatial scale differences effective scale differences were significantly larger for dDG compared to SM (*P* < 0.001) and to LoG (*P* < 0.034) and for LoG compared to SM (*P* < 0.032).

**Figure 7:**
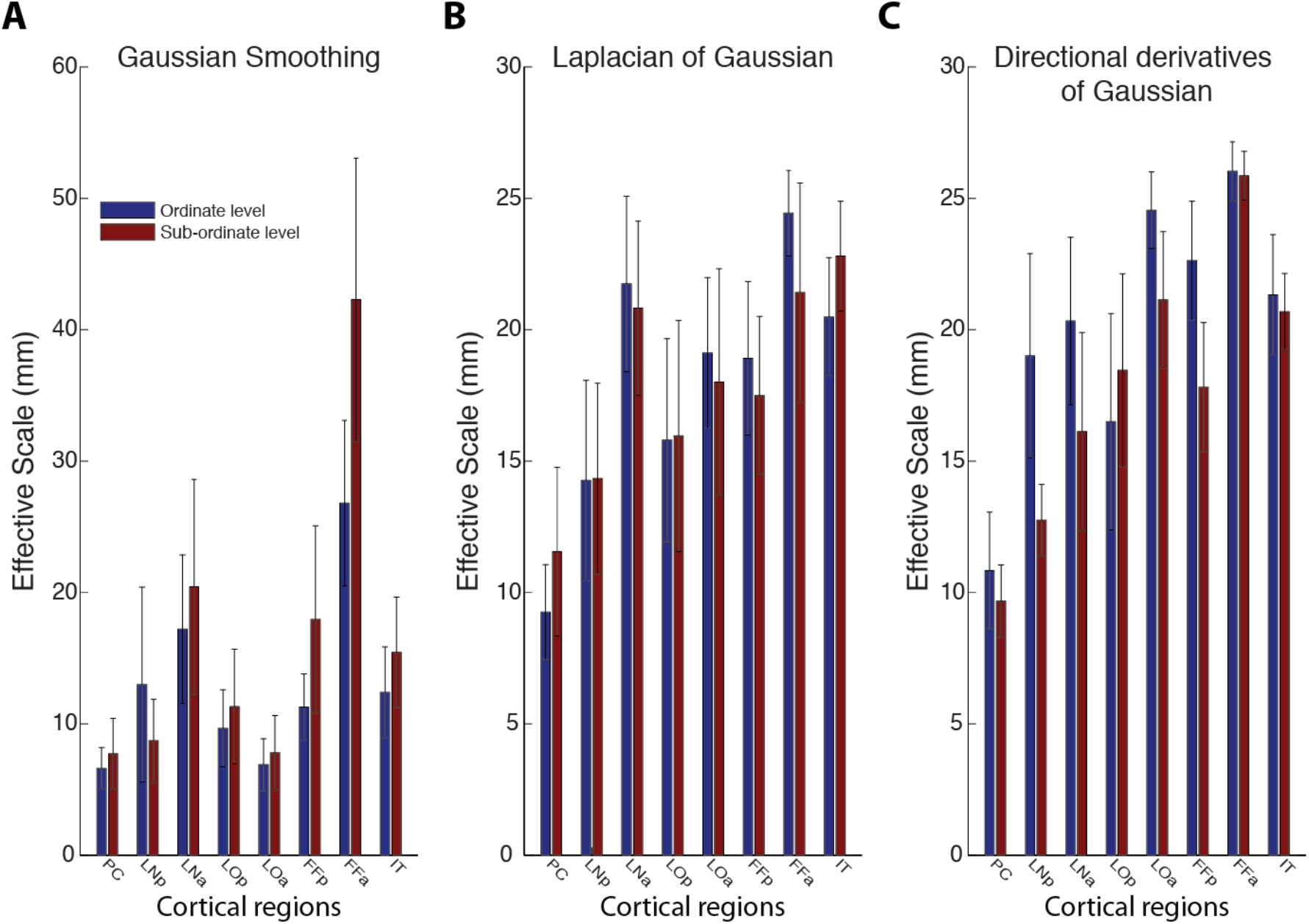
Effective scales at which the maximal *F*-statistics of ROIs is maximal for sub-ordinate and ordinate level distinctions among visual objects for (**A**) SM, (**B**) LoG and (**C**) dDG filtering. Only dDG, but neither SM nor LoG revealed a significant difference between categorical levels. Further, the effective scale at which information was encoded in ventral visual cortex increased with a posterior to interior gradient. Error bars indicate the standard error across subjects. Regions are ordered to approximately reflect the hierarchy in ventral visual cortex from posterior to anterior.

Together, these results show that dDG resolves differences in the spatial scale at which information is encoded in cortex where other methods fail, demonstrating the improved resolution of spatial scale of the dDG approach.

#### 2.3.3 The effective scale at which information is encoded in ventral visual cortex increases with a posterior-to-interior gradient

Visual inspection of the information map in Fig. 4 had suggested that the spatial scale at which information is encoded in ventral visual cortex might increase from posterior to anterior. The ROI analysis reinforced this observation (Fig. 7) the spatial scale at which classification was maximal at both the sub- and the supra-ordinate level increased along the processing path of the ventral visual stream from posterior to anterior

We thus quantified this observation by calculating Kendall’s tau rank correlation between the preferred scales and the ordinate position of the ROIs on the posterior-to-anterior axis of ventral visual cortex (ordered as the x-axis in Fig. 7). All filtering methods showed a positive correlation for both sub-ordinate and super-ordinate information encoding (Table 4). This result was ascertained statistically by one sided t-tests, revealing significant results for both levels of abstraction and all filtering methods (Table 4, all *P* < 0.05, FDR-corrected). Together, our results demonstrate a gradual increase in the spatial scale at which discriminant information is encoded along the cortical sheet of ventral visual cortex.

**Table 4:** Kendall rank correlation of scales across regions in ventral visual pathway at the basic and the sub-ordinate level.

|  | Ordinate level |  |  | Sub-ordinate level |  |  |
| --- | --- | --- | --- | --- | --- | --- |
|  | SM | LoG | dDG | SM | LoG | dDG |
| **Kendall's tau** | 0.2354 | 0.3000 | 0.2872 | 0.2635 | 0.2482 | 0.4215 |
| **p-value** | 0.0013 | <0.0001 | <0.0001 | 0.0003 | 0.0007 | <0.0001 |

## 3 Discussion

### 3.1 Summary

Here we present a novel analysis to determine the spatial scale and direction of activation patterns on the cortical sheet. Using an efficient algorithm for accurately computing Gaussian smoothing on cortical surfaces and discrete differential operators, we constructed wavelet-like bandpass filters with directionality and steerability for scale-specific analysis of cortical activity measurements. Evaluating the algorithm through modelling, we found increased precision compared to previous approaches. Applying the analysis to an fMRI data set of visual activation during object vision, we found that our analysis improved detection of discriminative information between experimental conditions, and provided novel insight into the cortical representations of objects: the spatial scale at which objects information is preferentially encoded depends on the level of categorization, and increase along the ventral visual pathway.

### 3.2 Smoothing and bandpass filtering on the irregular cortical sheet

#### 3.2.1 All algorithms for Gaussian smoothing are related, but differ in precision and complexity

What is the algorithmic nature of the proposed smoothing operator here, and how does it relate to the approaches compared? Note that all algorithms for Gaussian smoothing on the surface evaluated here can be formulated as the solution of the diffusion equation (1). They differ merely the choice of the discrete Laplace operator Δ or the respective heat kernel *e*^−*t*Δ^*δ*(*x*), and the numerical algorithm for implementing its application to the initial condition or input function *f*_0_.

In particular, iterative averaging (Hagler et al., 2006) is a linear approximation of the exponential operator applied to the input function, with the choice of normalized graph Laplacian (for proof see Supplementary Text II). While being the most popular choice for a smoothing operator, and one to two orders of magnitude *smaller* than the matrix exponential algorithm in computational complexity, the trade-off is inhomogeneity of smoothness and geometric deviation from Gaussian kernel. Thus, for detailed analyses of the spatial scale on irregular meshes a more sophisticated geometric discretization of the underlying Laplacian operator – as used here– is to be preferred.

#### 3.2.2 Choice of implementation of the exponential of Laplacian

In our approach, we adopt the geometrical Laplacian (4) and matrix exponential algorithm (Sidje, 1998) to implement the exponential of Laplacian. An alternative would have been to compute the exponential of Laplacian by explicitly solving the general eigen-decomposition of Δ (Seo et al. 2010). However, in practice the eigen-decomposition would have to be truncated and thus likely suffer from the rippling effects of spectral truncation and very high computational cost for explicit eigen-decomposition. Our approach avoids both of these shortcomings, improving both approximation precision and computational efficiency.

#### 3.2.3 The advantage and caveats of implementing bandpass filters by differential of smoothing

It is common practice in computer vision to implement isotropic bandpass filters like LoG by difference of Gaussians (DoG, Marr and Hildreth, 1980). Here, however, we instead adopted a direct approach to compute bandpass filtering by exploiting the differential property of convolution for two reasons. First, it is computationally more efficient when large support of filters is wanted, as it avoids calculating a much (typically 1.6-2x) larger Gaussian smoothing for DoG. Second, and more importantly, it allows combining first-order partial differential operators with the smoothed function to construct directional filters of derivatives of Gaussian. Note also that on a domain lacking a properly defined Fourier transform, such as an irregular mesh, multidimensional derivative filters cannot be designed directly as in Simoncelli (1994).

However, our approach has the caveat that precision relies heavily on the approximation quality of the discrete differential operator. Particularly, higher order partial differential operators cannot be constructed straightforwardly by recursive application of first-order partial differential operators, as differential of gradient vector field would have to deal with parallel transportation on the surface.

### 3.3 Effective filter size and effective spatial scale need to be assessed carefully

#### 3.3.1 The effective scale of results should be distinguished from the effective size of filters applied

Most previous studies analyzing fMRI data at multiple spatial scales relied on filter size as an indicator of the spatial scale of cortical patterns assessed (Swisher et al., 2010; Brants et al., 2011; Misaki et al., 2013). That is, they equated the effective scale of results with the effective size of the filters applied. Contrary to the appealing intuition underlying this interpretation, we argue that the effective scale of results needs to be determined independently by estimating the smoothness of residuals based on random field theory (Hagler et al., 2006).

The rationale behind this argument is straightforward: Whereas the effective filter size is estimated by using spatially independent Gaussian random noise as input functions, in neuroimaging data intrinsic spatial correlations are omnipresent at multiple scales due to various physiological and physical sources during the imaging procedure, and contribute to a noticeable increase of the effective scale of residuals compared to the effective filter size.

One particular observation in this regard is that bandpass filters have smaller effective sizes than smoothing filters of corresponding size. This might appear counter-intuitive at first sight, as the application of a discrete differential operator to a smoothing kernel should rather *increase* than *decrease* the support of the actual filter. However, a bandpass filter from a Gaussian family can be thought as a superimposition of its positive and negative parts, each of which has a support slightly bigger than half of the smoothing kernel. When calculated by RFT-based smoothness estimation, the effective filter size of such filters would approximately be the same as that of the parts (see the almost fixed ratio between the slopes in Fig. 4).

In sum, particular care needs to be taken when estimating the effective scale of results from neuroimaging data.

### 3.4 The role of bandpass filtering and steerability of the filters

#### 3.4.1 Steerability is necessary to fully characterize the scale property of discriminant information in cortex

Our results indicate that bandpass filters play an important role in characterizing the spatial scale properties of discriminant information encoded in cortex. The application of LoG and dDG not only showed an improved performance in discriminant analysis, but also revealed a systematic *increase* of scale along the ventral visual pathway. Concerning the further differentiation of LoG and dDG, the difference of characteristic scale between ordinate and subordinate categorization was only significant when dDG was applied, not LoG. This suggests that the improved characterization is more likely due to the steerability of the directional filters, rather than the bandpassing nature of these filters.

How is superior ability to detect information to be explained? Looking at steerable bandpass filters from different perspectives elucidates this issue. From the perspective of geometry, the optimal linear combination of directional derivative filters, as computed by the multivariate analysis in SurfStat, indicates a local direction along which the steepest change is statistically detected. From the perspective of wavelet analysis, steerable wavelets can be regarded as a special kind of matching pursuits (Bergeaud & Mallat, 1994), which achieve an optimal representation of the underlying discriminant information pattern in the space spanned by these wavelets. Finally, we may take the perspective of multivariate pattern analysis (MVPA) while changing the level of regularization. Spatial filters with specific shape may be considered as MVPA with very strong regularization. The strongest regularization, as in Gaussian smoothing kernels, permits only non-negative coefficients. Relaxing the regularization, such as LoG does by permitting negative coefficients and steerable filters with additional linear weights, allows better model fits. Interestingly, the very small number of parameters makes this approach far *less* likely to overfit than other common approaches of MVPA.

### 3.5 Implications for the understanding of the functional organization of ventral visual cortex

#### 3.5.1 Information differentiating objects at different levels of categorization is preferentially decodable at different scales

Our finding that discriminative information for ordinate categories is decodable preferentially at a coarser scale than that for sub-ordinate categories concurs with previous studies, both using fMRI in humans end electrophysiology in monkey (Tanaka et al., 2003; Op de Beeck et al., 2008; Brants et al., 2011). This further strengthens the idea that there is an ordered relationship between the topography of high-level ventral visual cortex and the hierarchy of visual object knowledge.

Note that the spatial scales reported here are much coarser than recently reported by joint analyses of neurophysiological and brain imaging data (Issa et al., 2013) in monkey. We believe that this discrepancy can be explained by the limited resolution of fMRI measurement investigated here, and that due to low SNR the analysis is most sensitive when pooling over a large number of voxels, and thus large spatial scales. Future studies, using ultra-high field fMRI and higher spatial resolutions will be necessary to resolve this open issue.

#### 3.5.2 Differences in preferential scale at which information is encoded across regions suggests different representational schemes

We observed an increase in the preferential scale at which object categorical information was decodable in regions along the ventral visual stream. This indicates a systematic change in functional organization at different stages of object processing hierarchy. The relatively fine scale in early visual cortex (e.g., PC, ~10mm; LNp, ~15mm) suggests a fine-tuned, retinotopically local encoding of similar object features in small cortical patches. In contrast, the relatively coarse scale in down-stream regions (e.g., LOa, >20mm; FFp, ~20mm) points to global and categorical organizing principles, such as gradients or topological maps indicating category (Grill-Spector & Weiner, 2014).

Our results inform about the nature of visual representations beyond the mere spatial scale in two ways. First, we observed that bandpass filtering outperforms *any* size of smoothing in determining the most discriminative information. This speaks against the idea that discriminant information is encoded in simple activated blobs such as inherent in the idea of univariate analysis of fMRI data, but is rather represented in inherent patterning with both positive and negative values, coupled geometrically over the cortical space. Second, we found that in discriminant analysis steerable filters outperformed symmetric filters across all regions and scales. This suggests that an intrinsic geometry in such patterning exists throughout from fine scale in clustering structures in early visual regions, to large scale topological map-like organization of high-level ventral visual cortex.

Future experiments investigating the detailed nature of representations of visual attributes other than object identity are necessary to establish the generality of these observations, and might benefit from the analysis framework proposed here.

### 3.6 SUMMARY

Together, our results indicate that the proposed analysis of activation patterns in scale and direction to be particularly suited to assess and detect scale specific information encoded by the cortical activity patterns, promising further insight into the topography of cortical functioning in the human brain.

## 4 Acknowledgements

This work was funded by the Bernstein Computational Neuroscience Program of the German Federal Ministry of Education and Research BMBF Grant 01GQ0411, the Excellence Initiative of the German Federal Ministry of Education and Research DFG Grants GSC86/1-2009, KFO247, HA 5336/1-1 and JA 945/3-1 / SL 185/1-1, and the Emmy Noether award CI 241-1/1.

## References

Bergeaud, F., & Mallat, S. (1994). Matching pursuit of images. In Time-Frequency and Time-Scale Analysis, 1994. Proceedings of the IEEE-SP International Symposium on (pp. 330–333). IEEE.

Biyikoglu, T., Leydold, J., & Stadler, P. F. (2007). Laplacian eigenvectors of graphs. Springer Lecture notes in mathematics, 1915.

Brants M, Baeck A, Wagemans J, Op de Beeck HP. (2011). Multiple scales of organization for object selectivity in ventral visual cortex. NeuroImage, 56(3), 1372–1381.

Brett, M., Johnsrude, I. S., & Owen, A. M. (2002). The problem of functional localization in the human brain. Nature reviews neuroscience, 3(3), 243–249.

Chaimow, D., Yacoub, E., Ugurbil, K., & Shmuel, A. (2011). Modeling and analysis of mechanisms underlying fMRI-based decoding of information conveyed in cortical columns. Neuroimage, 56(2), 627–642.

Chen, Y., Namburi, P., Elliott, L. T., Heinzle, J., Soon, C. S., Chee, M. W., & Haynes, J. D. (2011). Cortical surface-based searchlight decoding. Neuroimage, 56(2), 582–592.

Chung, M. K., Robbins, S. M., Dalton, K. M., Davidson, R. J., Alexander, A. L., & Evans, A. C. (2005). Cortical thickness analysis in autism with heat kernel smoothing. NeuroImage, 25(4), 1256–1265.

Cichy, R. M., Chen, Y., & Haynes, J. D. (2011). Encoding the identity and location of objects in human LOC. Neuroimage, 54(3), 2297–2307.

Dale, A. M., Fischl, B., & Sereno, M. I. (1999). Cortical surface-based analysis: I. Segmentation and surface reconstruction. Neuroimage, 9(2), 179–194.

Daubechies, I. (1990). The wavelet transform, time-frequency localization and signal analysis. Information Theory, IEEE Transactions on, 36(5), 961–1005.

Fischl, B., van der Kouwe, A., Destrieux, C., Halgren, E., Ségonne, F., Salat, D. H., Busa, E., Seidman, L. J., Goldstein, J., Kennedy, D., Caviness, V., Makris, N., Rosen, B. & Dale, A. M. (2004). Automatically parcellating the human cerebral cortex. Cerebral cortex, 14(1), 11–22.

Fischl, B., Sereno, M. I., & Dale, A. M. (1999). Cortical surface-based analysis: II: Inflation, flattening, and a surface-based coordinate system. Neuroimage, 9(2), 195–207.

Freeman, W. T. and Adelson, E. H. (1991). The design and use of steerable filters. IEEE Transactions on Pattern Analysis and Machine Intelligence, 13(9):891–906.

Freeman, J., Brouwer, G. J., Heeger, D. J., & Merriam, E. P. (2011). Orientation decoding depends on maps, not columns. The Journal of Neuroscience, 31(13), 4792–4804.

Freeman, J., Heeger, D. J., & Merriam, E. P. (2013). Coarse-scale biases for spirals and orientation in human visual cortex. The Journal of Neuroscience, 33(50), 19695–19703.

Goesaert, E., & de Beeck, H. P. O. (2010). Continuous mapping of the cortical object vision pathway using traveling waves in object space. Neuroimage, 49(4), 3248–3256.

Grill-Spector, K., & Weiner, K. S. (2014). The functional architecture of the ventral temporal cortex and its role in categorization. Nature Reviews Neuroscience, 15(8), 536–548.

Grinvald, A., Lieke, E., Frostig, R. D., Gilbert, C. D., & Wiesel, T. N. (1986). Functional architecture of cortex revealed by optical imaging of intrinsic signals. Nature, 324(6095), 361–364.

Haynes, J. D., & Rees, G. (2005). Predicting the orientation of invisible stimuli from activity in human primary visual cortex. Nature neuroscience, 5(5), 686–691.

Hagler, D. J., Saygin, A. P., & Sereno, M. I. (2006). Smoothing and cluster thresholding for cortical surface-based group analysis of fMRI data. Neuroimage, 33(4), 1093–1103.

Haxby, J. V., Gobbini, M. I., Furey, M. L., Ishai, A., Schouten, J. L., & Pietrini, P. (2001). Distributed and overlapping representations of faces and objects in ventral temporal cortex. Science, 293(5539), 2425–2430.

Hubel, D. H., & Wiesel, T. N. (1963). Shape and arrangement of columns in cat’s striate cortex. The Journal of physiology, 165(3), 559–568.

Issa, E. B., Papanastassiou, A. M., & DiCarlo, J. J. (2013). Large-scale, high-resolution neurophysiological maps underlying fMRI of macaque temporal lobe. The Journal of Neuroscience, 33(38), 15207–15219.

Kamitani, Y., & Tong, F. (2005). Decoding the visual and subjective contents of the human brain. Nature neuroscience, 5(5), 679–685.

Koenderink, J. J. (1984). The structure of images. Biological cybernetics, 50(5), 363–370.

Kanwisher, N., & Yovel, G. (2006). The fusiform face area: a cortical region specialized for the perception of faces. Philosophical Transactions of the Royal Society B: Biological Sciences, 361(1476), 2109–2128.

Logothetis, N. K., & Wandell, B. A. (2004). Interpreting the BOLD signal. Annu. Rev. Physiol., 66, 735–769.

Maldonado, P. E., Gödecke, I., Gray, C. M., & Bonhoeffer, T. (1997). Orientation selectivity in pinwheel centers in cat striate cortex. Science, 276(5318), 1551–1555.

Mallat, S. (2008). A wavelet tour of signal processing: the sparse way. Academic press.

Marr, D., & Hildreth, E. (1980). Theory of edge detection. Proceedings of the Royal Society of London. Series B. Biological Sciences, 207(1167), 187–217.

Meyer, M., Desbrun, M., Schröder, P., & Barr, A. H. (2003). Discrete differential-geometry operators for triangulated 2-manifolds. Visualization and mathematics III 35–57. Springer.

Misaki, M., Luh, W. M., & Bandettini, P. A. (2013). The effect of spatial smoothing on fMRI decoding of columnar-level organization with linear support vector machine. Journal of neuroscience methods, 212(2), 355–361.

Op de Beeck, H. P., DiCarlo, J. J., Goense, J. B., Grill-Spector, K., Papanastassiou, A., Tanifuji, M., & Tsao, D. Y. (2008). Fine-scale spatial organization of face and object selectivity in the temporal lobe: do functional magnetic resonance imaging, optical imaging, and electrophysiology agree?. The Journal of Neuroscience, 25(46), 11796–11801.

Portilla, J., & Simoncelli, E. P. (2000). A parametric texture model based on joint statistics of complex wavelet coefficients. International Journal of Computer Vision, 40(1), 49–70.

Ramírez, F. M., Cichy, R. M., Allefeld, C., & Haynes, J. D. (2014). The Neural Code for Face Orientation in the Human Fusiform Face Area. The Journal of Neuroscience, 34(36), 12155–12167.

Reuter, M., Biasotti, S., Giorgi, D., Patanè, G., & Spagnuolo, M. (2009). Discrete Laplace–Beltrami operators for shape analysis and segmentation. Computers & Graphics, 33(3), 381–390.

Rust, N. C., & DiCarlo, J. J. (2010). Selectivity and tolerance (“invariance”) both increase as visual information propagates from cortical area V4 to IT. The Journal of Neuroscience, 30(39), 12978–12995.

Seo, S., Chung, M. K., & Vorperian, H. K. (2010). Heat kernel smoothing using Laplace-Beltrami eigenfunctions. In Medical Image Computing and Computer-Assisted Intervention–MICCAI 2010 (pp. 505–512). Springer Berlin Heidelberg.

Shmuel, A., Chaimow, D., Raddatz, G., Ugurbil, K., & Yacoub, E. (2010). Mechanisms underlying decoding at 7 T: ocular dominance columns, broad structures, and macroscopic blood vessels in V1 convey information on the stimulated eye. Neuroimage, 49(3), 1957–1964.

Simoncelli, E. P., & Freeman, W. T. (1995). The steerable pyramid: A flexible architecture for multi-scale derivative computation. International Conference on Image Processing, (Vol. 3, 3444–3444). IEEE Computer Society.

Sidje, R. B. (1998). Expokit: a software package for computing matrix exponentials. ACM Transactions on Mathematical Software (TOMS), 24(1), 130–156.

Swisher, J. D., Gatenby, J. C., Gore, J. C., Wolfe, B. A., Moon, C. H., Kim, S. G., & Tong, F. (2010). Multiscale pattern analysis of orientation-selective activity in the primary visual cortex. The Journal of Neuroscience, 30(1), 325–330.

Unser, M., Chenouard, N., & Van De Ville, D. (2011). Steerable Pyramids and Tight Wavelet Frames. IEEE Transactions on Image Processing, 20(10), 2705–2721.

Van Essen, D. C., Drury, H. A., Joshi, S., & Miller, M. I. (1998). Functional and structural mapping of human cerebral cortex: solutions are in the surfaces. Proceedings of the National Academy of Sciences, 95(3), 788–795.

Van Essen, D. C., Lewis, J. W., Drury, H. A., Hadjikhani, N., Tootell, R. B., Bakircioglu, M., & Miller, M. I. (2001). Mapping visual cortex in monkeys and humans using surface-based atlases. Vision research, 41(10), 1359–1378.

Van Essen, David C., and Donna L. Dierker. "Surface-based and probabilistic atlases of primate cerebral cortex." Neuron 56.2 (2007): 209–225.

Wang, B., Hikino, Y., Imajyo, S., Ohno, S., Kanazawa, S., & Wu, J. (2012). Effect of spatial smoothing on regions of interested analysis basing on general linear model. International Conference on Mechatronics and Automation (ICMA) (1399–1404). IEEE.

Worsley, K. J., Jonathan E. Taylor, F. Carbonell, M. K. Chung, E. Duerden, B. Bernhardt, O. Lyttelton, M. Boucher, and A. C. Evans. (2009) SurfStat: A Matlab toolbox for the statistical analysis of univariate and multivariate surface and volumetric data using linear mixed effects models and random field theory. Neuroimage (47). (software package available at www.math.mcgill.ca/keith/surfstat)

